# Baseline power of theta oscillations predicts mystical-type experiences induced by DMT

**DOI:** 10.1101/2021.03.11.434994

**Authors:** Enzo Tagliazucchi, Federico Zamberlan, Federico Cavanna, Laura de la Fuente, Celeste Romero, Yonatan Sanz Perl, Carla Pallavicini

## Abstract

N,N-Dimethyltryptamine (DMT) is a classic psychedelic capable of inducing short-lasting but profound changes in consciousness. As with other psychedelics, the experience induced by DMT strongly depends upon contextual factors, yet the neurobiological determinants of this variability remain unknown. We combined wireless electroencephalography and source imaging to map changes in neural oscillations elicited by inhaled DMT. Furthermore, we found that the power of frontal and temporal theta oscillations was inversely correlated with scales indexing feelings of unity and transcendence, which are an integral part of the phenomenology of mystical-type experiences. Finally, we established the robustness of these results using a machine learning model for regression trained and tested following a cross-validation procedure. Our results are consistent with the observation that the state of mind prior to consuming a psychedelic drug influences the ensuing subjective experience of the user. We also suggest that priming subjects to reduce their theta power before administration of a serotonergic psychedelic could enhance the likelihood of inducing mystical-type experiences, leading to sustained positive effects in well-being and improving the outcome of therapeutic interventions.

## Introduction

Serotonergic psychedelics (a subset of 5-HT_2A_ receptor agonists) are capable of inducing remarkable changes in perception, affect, self-awareness and cognition (Nichols, 2016). Some of the effects elicited by psychedelics are long-lasting, for instance, changes in personality traits and improvements in the symptoms of certain mental health conditions (Bouso et al., 2018; Aday et al., 2020; Nutt and Carhart-Harris, 2021). After decades of neglect, psychedelics have resurfaced into the mainstream of human neuroscience and psychiatry, a revolution spearheaded by neuroimaging studies in healthy volunteers (Dos Santos et al., 2016) and clinical research that supports the efficacy of these drugs for the treatment of depression (Carhart-Harris et al., 2016; Palhano-Fontes et al., 2019; Davis et al., 2020), substance abuse (Johnson et al., 2014), and existential anxiety (Grob et al., 2011; Griffiths et al., 2016), among other uses. These advances, combined with the excellent safety profile of all classic psychedelics (mescaline, psilocybin, lysergic acid diethylamide [LSD] and N,N-dimethyltryptamine [DMT]) (Johnson et al., 2008), resulted in a surge of scientific studies aimed to understand how psychedelics exert their effect on consciousness and how these effects relate to their therapeutic mechanism of action (Sessa, 2018).

The capacity of psychedelics to induce long-lasting psychological effects is related to the qualities of the subjective experience during the acute effects of the drug. When administered in a pleasant and supportive setting, psilocybin is capable of inducing profoundly meaningful experiences of deep personal significance, with a sustained positive effect in well-being and behavior (Griffiths et al., 2006; Griffiths et al., 2008). The subjective experience linked to these transformative properties of psychedelics has substantial commonalities with other non-ordinary states of consciousness, known in the previous literature as mystical experiences, conversion experiences, peak experiences, transcendental experiences, epiphanies, among other denominations (Barrett and Griffiths, 2017; Johnson et al., 2019). Because of these similarities, Griffiths and colleagues introduced the name “mystical-type” to refer to these experiences, and built upon previous work to create the Mystical Experience Questionnaire (MEQ30), with a total of 30 items leading to four subscale scores (mystical, positive mood, transcendence of time and space, and ineffability) (Barrett et al., 2015). Psilocybin-induced mystical-type experiences are dose dependent (Griffiths et al., 2011) and are related to the therapeutic effects of the drug when administered in patients suffering from depression (Roseman et al., 2018), end of life anxiety (Griffiths et al., 2016) and substance abuse disorders (Garcia-Romeu et al., 2014), thus prompting the need to understand how these experiences can be predicted and facilitated.

Considerable variability exists in the percentage of subjects whose report qualifies as a complete mystical-type experience, according to the criteria put forward by Griffiths and colleagues. These percentages range from 60% for multiple studies using psilocybin (Barrett and Griffiths, 2017; Johnson et al., 2019) to 10% for LSD (Liechti et al., 2017), and between 37% and 73% for intravenous or inhaled DMT (Griffiths et al., 2019; Pallavicini et al., 2020). This variability is likely related to contextual factors known as set and setting, i.e. the state of mind of the participant prior to the experience and the environment where the experience takes place, respectively (Carhart-Harris et al., 2018). For instance, Haijen and colleagues conducted a large online survey to predict the response to psychedelic drugs, finding that trait absorption and clear intentions are conducive to mystical-type experiences (Haijen et al., 2018). A recent meta-analysis identified other predictors of mystical-type experiences, such as age, apprehension, deservingness, spiritual motivations, surrender, acceptance and attachment anxiety (Aday et al., 2021).

Facilitating mystical-type experiences is only one among multiple challenges related to the optimization of set and setting with the purpose of maximizing the therapeutic gain of treatment with psychedelics. Most predictive variables explored to date were obtained from self-reported measures assessed using standardized questionnaires and meta-analyses (Studerus et al., 2012; Haijen et al., 2018; Aday et al., 2021). However, insofar the state of mind of the user is reflected in baseline measurements of brain activity, these measurements should also present predictive power concerning the ensuing psychedelic experience, thus informing the neurobiological mechanisms underlying the repertoire of possible responses to psychedelic drugs. Notably, the development of predictive models based on brain activity recordings remains heavily underexplored in comparison to those based on psychometric data.

We explored baseline EEG oscillations as predictors of self-reported subjective effects in a cohort of 35 subjects who inhaled DMT in freebase form in their preferred context of use. DMT is frequently consumed in ceremonial settings, where it is crystallized over non-psychoactive plant leaves and then inhaled after combustion (Cakic et al., 2010; Winstock et al., 2014), leading to intense but short-lasting subjective effects (Szara, 1956; Strassman et al., 1994; Shulgin and Shulgin, 1997). We chose to investigate DMT under natural conditions to encompass an ample range of sets and settings, an advantage when searching for contextual factors that are linked to specific drug-induced experiences (Carhart-Harris et al., 2018; Shamay-Tsoory et al., 2019). Our main result is a link between mystical-type experiences and baseline theta oscillations originating from frontal and temporal brain regions.

## Materials and methods

The data included in this manuscript was used in a previous publication, which can be referenced for further methodological details (Pallavicini et al., 2020).

### Participants

Thirty-five participants (7 females; 33.1 ± 6 years; 92.2 ± 201.4 previous experiences with ayahuasca; 3.6 ± 5.6 previous experiences with DMT alone) were recruited by word-of-mouth and social media advertising between May and December 2019.

Participants were required to have at least two previous experiences with ayahuasca or DMT, abstain from consuming psychoactive drugs at least 24 hours prior to the study, and to be willing to engage in their preferred use of DMT in the presence of research team members. Researchers did not provide DMT or other psychoactive compounds to the subjects, nor interacted with their use of the substance. All subjects were aged between 21 and 65 years, and pregnant women were excluded from the experiment.

Subjects were excluded due to reported past difficult experiences with psychedelics, and based on the results of a non-diagnostic psychiatric interview (SCID-CT; First, 2014) according to the guidelines by Johnson et al. (2008). For further details concerning the inclusion and exclusion criteria see Pallavicini et al. (2020).

This study was conducted in accordance with the Helsinki declaration and approved by the Committee for Research Ethics at the Jose Maria Ramos Mejia General Hospital (Buenos Aires, Argentina).

All research data associated with this manuscript is publicly available without restrictions (10.5281/zenodo.3992359)

### DMT administration

Subjects consumed DMT in their preferred context of use. After being fitted with the EEG cap, the subjects were instructed to keep their eyes closed, relax and maintain an upright position, avoiding head movement and jaw clenching to prevent muscle artifacts. Only four subjects self-administered DMT, all others were assisted by a facilitator. After receiving instructions by the facilitators, subjects inhaled the smokes and vapors resulting from the combustion of freebase DMT, in all cases recrystallized over non-psychoactive plant leaves. Facilitators withdrew the pipe when subjects either stopped inhaling and leaned back, or exhausted the contents of the pipe. The average load of the pipes was estimated by the participants or their facilitators at 40 mg freebase DMT, in all cases extracted from the root of Mimosa hostilis (also known as Mimosa tenuiflora or jurema) (Ott, 1999). The presence of DMT was verified in all samples by high performance liquid chromatography coupled to mass spectrometry for profiling and qualitative analysis.

### Psychometric questionnaires

Before the DMT experience, participants completed Spanish versions of the State Trait Anxiety Inventory (STAI trait) (Spielberger, 2010), and questions introduced in Table 3 of Haijen et al. (2018) to assess the self-reported adequateness of set, setting and intentions. After the DMT experience, participants completed the 5D altered states of consciousness scale (5D-ASC) (Studerus et al, 2010), the mystical experience questionnaire (MEQ30) (Barrett et al., 2015), the near-death experience scale (NDE) (Greyson, 1983), and a series of questions to assess the impact of set, setting and social interactions on the psychedelic experience (post-experience questionnaire, or “Post”). Immediately before and after the DMT experience, participants completed Spanish versions of the Big Five personality (BFI) test (John and Srivastava, 1999), STAI state (Spielberger, 2010), and Tellegen absorption scale (TAS) (Tellegen and Atkinson, 1974).

### EEG acquisition and pre-processing

EEG data were recorded with a 24-channel mobile system (mBrainTrain LLC, Belgrade, Serbia; http://www.mbraintrain.com/) attached to an elastic electrode cap (EASYCAP GmbH, Inning, Germany; http://www.easycap.de). Twenty-four Ag/AgCl electrodes were positioned at standard 10– 20 locations (Fp1, Fp2, Fz, F7, F8, FC1, FC2, Cz, C3, C4, T7, T8, CPz, CP1, CP2, CP5, CP6, TP9, TP10, Pz, P3, P4, O1, and O2). Reference and ground electrodes were placed at FCz and AFz sites. The wireless EEG DC amplifier (weight = 60 g; size = 82 × 51 × 12 mm; resolution = 24 bit; sampling rate = 500 Hz, 0–250 Hz pass-band) was attached to the back of the electrode cap (between electrodes O1 and O2) and sent digitized EEG data via Bluetooth to a Notebook held by a experimenter sitting behind the participant. Prior to the administration of DMT, baseline EEG activity was acquired with eyes open and closed (five minutes each). After the DMT was administered, EEG recordings started when subjects exhaled, and lasted until the subject indicated a return to baseline (6 ± 1.4 min).

EEG data was preprocessed using EEGLAB (https://sccn.ucsd.edu/eeglab/index.php) (Delorme and Makeig, 2004). Data was divided into 2 s epochs, then bandpass-filtered (1 – 90 Hz) and notch-filtered (47.5 – 52.5 Hz). Artifact-laden channels were first detected using EEGLAB automated methods. All channels were manually inspected before rejection (mean 30% rejected channels, max. 8 channels) and then interpolated using data from the surrounding channels. Epochs to be rejected were flagged automatically and in all cases removed after manual inspection (21.3 ± 13.7 epochs rejected per subject). Infomax independent component analysis (ICA) was then applied to data from each individual participant, and used to manually identify and remove artifactual components (2.7 ± 1.1 components removed). According to the previously defined criteria, 6 subjects were discarded from the subsequent EEG analysis due to an excessive number of rejected epochs and/or channels, resulting in 29 subjects for subsequent analysis.

### Source EEG power estimation

Source imaging analysis was performed using Brainstorm (https://neuroimage.usc.edu/brainstorm/) (Tadel et al., 2011). Pre-processed EEG epochs were imported and localized to a standard MRI volume divided by a 4 mm 3D grid. A 3-shell sphere model was used to compute the forward solutions, with adaptive integration and standard tissue conductivities. Sources were estimated by minimum norm imaging with unconstrained dipole orientations and identity noise covariance matrix, yielding a current density map normalized by the square root of the estimated noise variance at each location in the map, i.e. a z-score statistical map (dynamical Statistical Parametric Mapping [dSPM]) (Dale, 2000).

Band-specific power spectral density was computed using the Welch method followed by spectrum normalization and averaging across epochs. Spectral power was computed for the following frequency bands: delta (1–4 Hz), theta (4–8 Hz), alpha (8–12 Hz), beta (12–30 Hz), low gamma (30–40 Hz) and high gamma (40–70 Hz). Finally, source spectral power estimates were averaged across all grid locations within the cortical and subcortical regions defined in the Automated Anatomical Labeling (AAL) atlas, yielding 90 values per subject and frequency band (Tzourio-Mazoyer et al., 2002).

### Statistical analyses

Significant differences in regional mean spectral power between conditions were assessed using Wilcoxon’s nonparametric signed-rank test. The predictive power of baseline activity was evaluated using Spearman’s nonparametric rank correlation coefficient between regional spectral estimates during the eyes closed condition and the results of questionnaires (5D-ASC, NDE and MEQ30) obtained after the DMT condition.

The Benjamini-Hochberg procedure was followed to control for the false discovery rate, setting a rate of *α*=0.05 (Benjamini and Hochberg, 1995) Correlation p-values obtained for all frequency bands and questionnaire subscales were pooled together before the application of this procedure.

### Machine learning regression model

A random forest regressor (Liaw and Wiener, 2002) with 1000 estimators (implemented in scikit-learn, https://scikit-learn.org/) (Pedregosa et al., 2011) was trained using 50% of the samples to estimate the 5D-ASC Unity, MEQ30 Mystical and MEQ30 Transcendence scores in the remaining 50% (randomly selected for each independent instance of the classifier). Each sample consisted of the theta power source values at the 90 AAL regions for a participant during the eyes closed baseline. This procedure was repeated 1000 times for each questionnaire subscale with and without shuffling of the target scores. Finally, a p-value was constructed by counting the number of times the mean squared error of the randomized classifier was lower than that of the classifier trained using the non-shuffled scores, divided by the total number of iterations.

## Results

### Differences in source EEG power between DMT and baseline

We first compared the EEG source estimates for all frequency bands between the eyes closed and DMT conditions. The results of this analysis are shown in Fig. 1. We observed posterior increases in delta power under the acute effects of DMT, with similar (but less marked) changes for the theta band. For the alpha band we observed changes similar to those seen in the delta band, but of opposite sign (i.e. alpha power decreases in occipito-temporal regions under DMT). Beta power increased in posterior regions and decreased in prefrontal regions, while occipital, parietal and temporal gamma increases appeared under DMT. These results are consistent with those previously reported at the scalp level (Pallavicini et al., 2002), but with significant differences in more frequency bands (mainly theta and beta).

**Figure 1.**
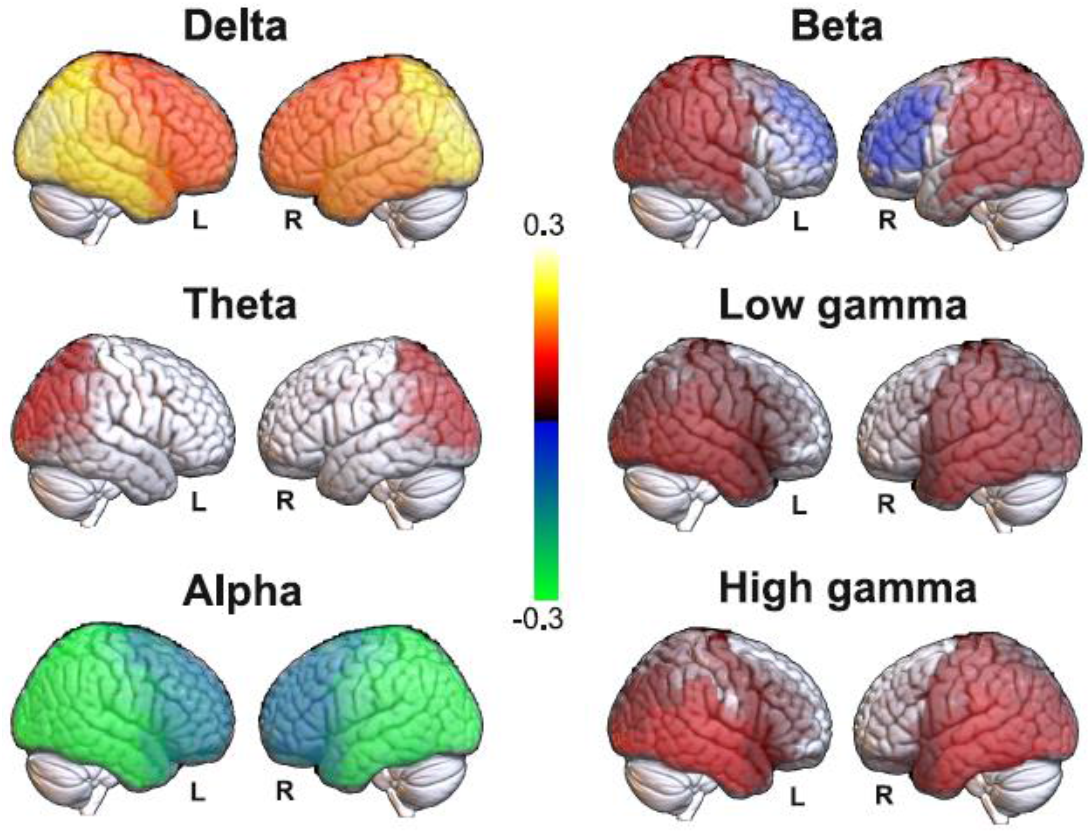
Significant differences in band-specific source EEG power between the eyes closed baseline and the DMT condition. Red-yellow and blue-green color scales indicate decreased and increased spectral power under DMT compared to the baseline, respectively.

### Correlations between baseline activity and questionnaire subscales

Next, we computed all correlations between regional spectral power estimates and questionnaire subscales. After FDR-correction for multiple comparisons, we only identified significant correlations for the theta band (see Fig. 2 for a matrix summarizing all correlations for this frequency band). These significant correlations corresponded to the following three subscales: 5D-ASC Experience of unity (“Unity”), MEQ30 Mystical experience (“Mystical”) and MEQ30 Transcendence of time and space (“Transcendence”).

**Figure 2.**
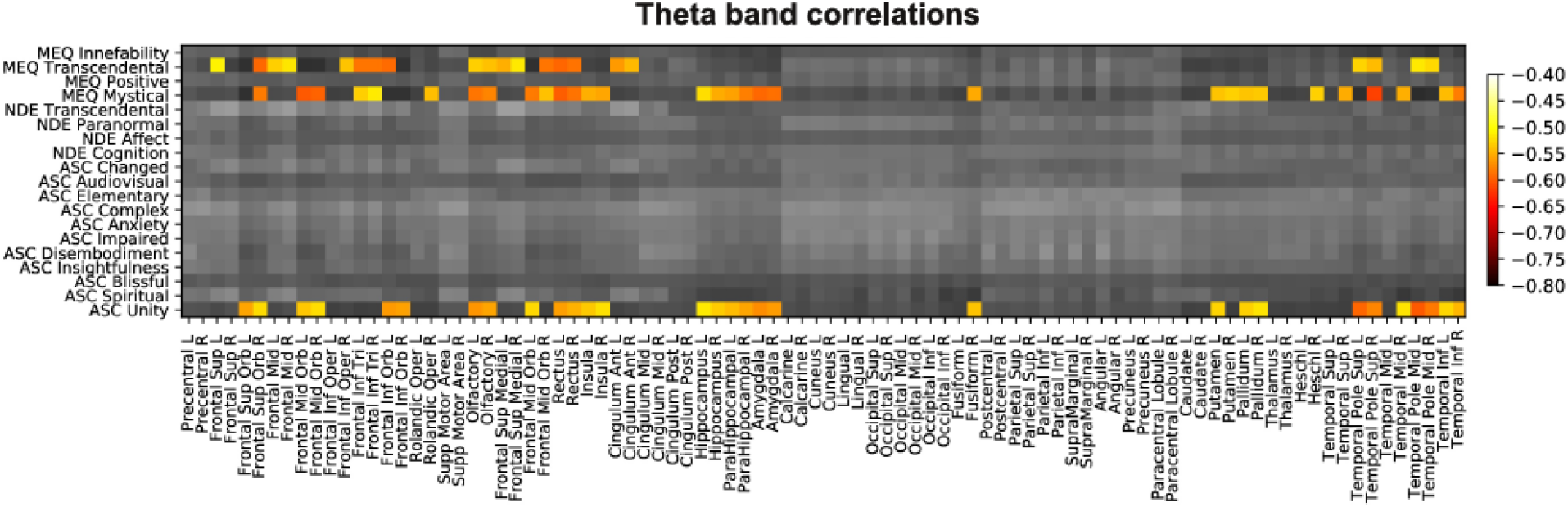
Matrix with all correlations between questionnaire subscales (y-axis) and source theta power computed for all AAL regions (x-axis). Significant correlations were in all cases negative and are indicated by colored entries.

Figure 3 presents the anatomical rendering and volumetric display of the AAL regions whose baseline theta power presented significant correlations with 5D-ASC Unity, MEQ30 Mystical and MEQ30 Transcendence. 5D-ASC Unity was inversely correlated with theta power at fronto-temporal regions, including the bilateral hippocampus and parahippocampal gyrus. A similar pattern of significant correlations was observed for MEQ30 Mystical, while negative correlations between theta power and MEQ30 Transcendence were distributed frontally and bilaterally, spanning the prefrontal cortex, the orbital part of the frontal lobe and the anterior cingulate cortex.

**Figure 3.**
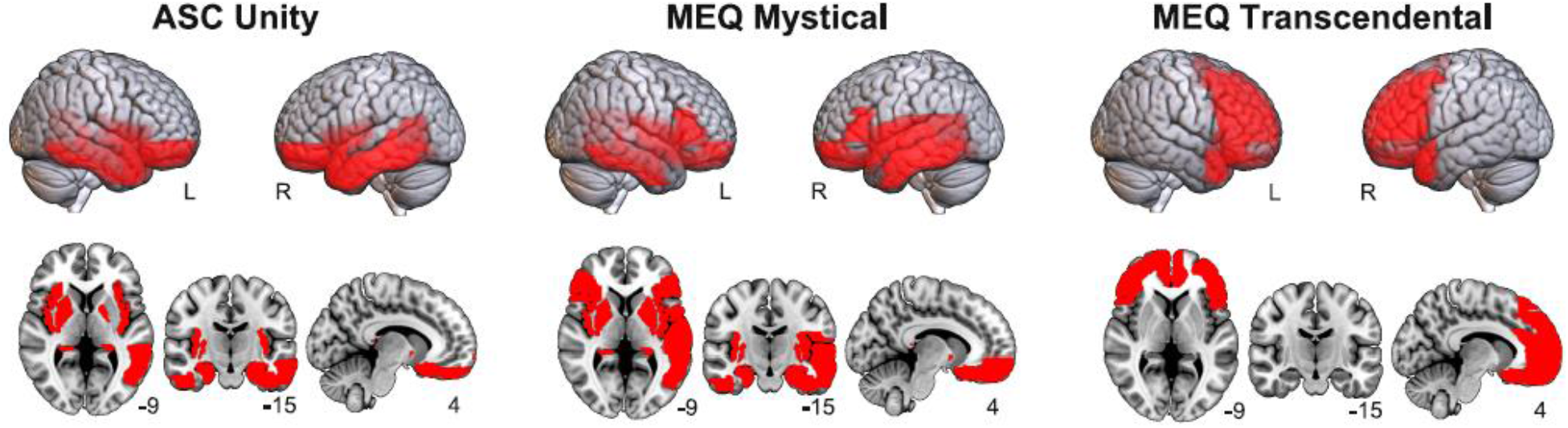
Anatomical rendering (top) and volumetric display (bottom) of AAL regions where significant negative correlations between theta power and questionnaire subscales (5D-ASC Unity, MEQ30 Mystical and MEQ30 Transcendental) were found.

### Prediction using a cross-validated machine learning regression model

Figure 4 shows the prediction of individual questionnaire subscale values based on the theta power localized at the 90 AAL regions, with the y-axis representing the predicted values (obtained using a random forest regression model) and the x-axis representing the actual values. In both cases, values were standardized to z-scores before visualization. As shown in the insets, the predicted values presented a medium to large correlation with the actual values; however, the model tended to underestimate the scores that were significantly smaller than the mean (i.e. largest negative values after conversion to z-scores).

**Figure 4.**
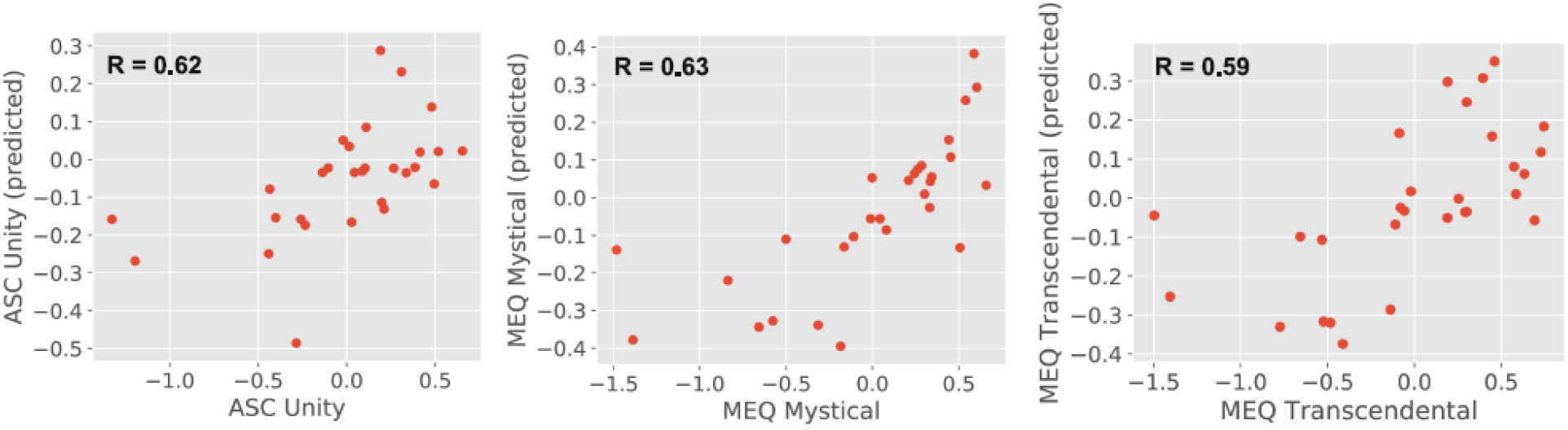
Predicted (random forest regression model) vs. actual questionnaire subscale values (in both cases standardised to z-scores) with the linear correlation coefficient between both included as an inset.

### Prediction based on psychometric data

We computed the correlation between 5D-ASC Unity, MEQ30 Mystical and MEQ30 Transcendence subitem scores, and baseline psychometric scores including personality traits (BFI), trait anxiety (STAI) and trait absorption (TAS). Consistent with a previous survey study (Haijen et al., 2018), we only found significant correlations with trait absorption (R=0.43 for 5D-ASC Unity and R=0.51 for MEQ30 Mystical).

## Discussion

We investigated how the acute effects of DMT modified source-localized EEG spectral power, as well as the relationship between baseline spectral power fluctuations and the conscious experience reported by the subjects. Our recordings were obtained in natural settings, i.e. the contexts preferred by the participants for their use of DMT. While this approach is limited in many ways compared to a double-blind placebo-controlled design, it also presents distinct advantages, mainly related to an ample range of settings. These advantages are especially important when attempting to predict the response to DMT from baseline brain activity measurements (Carhart-Harris et al., 2018; Shamay-Tsoory et al., 2019). We followed a novel approach, diverging from previous efforts to predict the effects of psychedelics from questionnaires or from natural language descriptions of set and setting (Studerus et al., 2012; Haijen et al., 2018; Aday et al., 2021). Since the mindset of the participants should be reflected on their baseline neural oscillations, we hypothesized that band-specific EEG power could predict some of the subjective effects reported by them after the DMT experience.

Our results at the source level are consistent with a previous analysis of this data (Pallavicini et al., 2020) and with an independent report of EEG changes elicited by intravenous DMT administration (Timmermann et al., 2019). The most salient results obtained from the contrast of DMT vs. eyes closed comprise increased delta power and decreased alpha power, in both cases localized to posterior brain regions, replicating previous analyses. Also, we found that DMT increased the power of gamma oscillations in occipital and temporal regions; a similar increase of oscillations in the gamma range was previously reported by our group, as well as by others investigating orally administered DMT in the form of ayahuasca (Schenberg et al., 2015). However, we also found new significant changes in the theta and beta frequency bands, suggesting that mixed sources registered at the scalp could hinder the level of significance in the comparison of DMT vs. eyes closed.

Theta oscillations have been linked to multiple cognitive and perceptual processes in the healthy human brain, some of them subserving mind-wandering (Klimesch, 1999; Sauseng et al., 2005; Demiralp et al., 2007; Sauseng et al., 2010). We speculate that these functions could be detrimental for a state of mind conducive to mystical-type experiences, an observation supported by the facilitation of these experiences by mindfulness training (Smigielski et al., 2010). Increased theta power has been reported during mind-wandering episodes, especially when divided by beta power to form the beta/theta ratio (van Son et al., 2019a, van Son et al., 2019b). Theta oscillations have been linked to episodes of self-assessed (i.e. subjective) drifts of attention during a breath counting task (Braboszcz and Delorme, 2011), as well as to behaviorally indexed mind-wandering (Qin et al., 2011). Dissociative absorption, a trait predictive of psychedelic-induced mystical-type experiences, correlates inversely with the power of theta rhythms originating within the temporal lobes (Soffer-Dudek et al., 2019). A combined EEG-fMRI study established that intrusions of self-generated and internally-oriented thought processes are characterized by reduced default mode network activity and increased power of theta oscillations, among other frequency bands (Groot et al., 2021). Anxiety prior to the DMT experience could hinder the likelihood of undergoing a mystical-type experience, which is consistent with the reported link between anxious rumination and increased theta power localized to the anterior cingulate cortex (Andersen et al., 2009).

Oscillations in the theta band are thought to reflect top-down influences related to working memory and memory retrieval (Klimesch, 1999; Sauseng et al., 2005; Demiralp et al., 2007; Sauseng et al., 2010), and are linked to the activation of autobiographical memories (Steinvorth et al., 2010) as well as to mental time travel (Lavallee and Persinger, 2010) and, more generally, to the level of mental workload (So et al., 2017). While these aspects of theta oscillations could be related to mind-wandering, multiple reports show enhanced theta power in mindfulness meditators, as well as in meditators following other contemplative traditions (Lagopoulos et al., 2009; Xue et al., 2014; Lomas et al., 2015; Harne and Hiwale, 2018). Furthermore, frontal theta oscillations are inversely correlated with default mode activity, suggesting that these oscillations signal episodes of sustained attention (Scheeringa et al., 2008). These apparent contradictions could stem from the lack of a unique source for the theta rhythm, with different sources reflecting equally distinct cognitive roles, and from the possibility of rhythms of the same frequency but different functional roles emerging from the same cortical regions (Zuure et al., 2020). Since we did not assess the prevalence of mind-wandering episodes prior to the DMT experience, we lack subjective or behavioral validation for the proposed disruptive role of theta oscillations concerning mystical-type experiences. However, we note that high absorption is predictive of these experiences (as previously shown and confirmed in our analysis) and at the same time indicative of deep and effortless concentration (Berkovich-Ohana and Glicksohn, 2017) and inversely correlated with EEG theta power (Soffer-Dudek et al., 2019).

Our analysis failed to predict other aspects of the psychedelic experience, such as perceptual, affective or cognitive alterations. Haijen and colleagues showed that the intensity of visual effects could be predicted by absorption, dose, and having clear intentions for the experience (Haijen et al., 2018). However, other studies suggest that predicting perceptual modifications from non-pharmacological variables is more difficult than predicting mystical-type experiences (Studerus et al., 2012). A recent meta-analysis established that baseline psychometric questionnaires are mostly predictive of affective or mystical-type experience, with only sparse significant associations of baseline variables with perceptual or cognitive modifications (Aday et al., 2021). We can speculate that individual characteristics beyond the resolution of EEG could underlie the variability in other dimensions of the psychedelic experience, for instance, differences in the cortical density of serotonin 5-HT2A receptors. Future studies should attempt to predict the response to psychedelics from more exhaustive neuroimaging recordings, including functional, anatomical, and neurochemical data.

Our study is limited by lack of a placebo condition to control for expectancy effects, as well as by limited information concerning the administered dose of DMT. These limitations are inherent to studies conducted in natural settings and thus future research should attempt to reproduce our findings following a randomized double-blind placebo-controlled design. However, we note that predicting how contextual factors influence the acute effects of psychedelics might be difficult if variables related to set and setting are uniform across participants. In this sense, a certain degree of contextual heterogeneity (as in the case of naturalistic studies) could represent an advantage instead of a limitation.

In summary, our study represents a first step in the direction of predicting the acute effects of psychedelics from baseline neurophysiological recordings. Although debated, the therapeutic properties of psychedelics seem to depend upon the state of consciousness manifested during the experience itself. As more clinical research is conducted and more data becomes available, we expect that certain profiles of subjective effects will be associated with the improvement of patients after treatment with psychedelics. Ultimately, the field will face the problem of how to engineer these desirable subjective effects, a problem that will require a systematic exploration of the response to psychedelic drugs from baseline psychological and neurophysiological measurements.

## Acknowledgments

We thank Gonzalo Sierra for his continuous support of this study. The authors also acknowledge Toyoko LLC for granting cloud computing services. This work was supported by funding from Agencia Nacional De Promocion Cientifica y Tecnologica (Argentina), grant PICT-2018-03103. The authors declare no conflict of interest.

## Notes

https://zenodo.org/record/3992359

### Competing Interest Statement

The authors have declared no competing interest.

